# A Robust Isotope Ratio LC-MS/MS Workflow for High-Throughput Metabolic Profiling of Bacteria

**DOI:** 10.1101/2025.08.08.669109

**Authors:** Catarina Rocha, Sergi Pujol Pinto, Sheila Ingemann Jensen, Lars Keld Nielsen, Stefano Donati

**Affiliations:** The Novo Nordisk Foundation Center for Biosustainability, Technical University of Denmark, Kongens Lyngby, Denmark; Australian Institute for Bioengineering and Nanotechnology (AIBN), The University of Queensland, Brisbane, Queensland, Australia; ARcCentre of Excellence in SyntheticBiology, The University of Queensland, Queensland, Australia

**Keywords:** Quantitative intracellular metabolomics, Ion-pairing LC IDMS, ^13^C-labeled IS, Cross-species metabolomics, microbiology, systems biology

## Abstract

Stable isotope dilution mass spectrometry (IDMS) has become a cornerstone of quantitative metabolomics, enabling accurate intracellular metabolite quantification across a range of biological systems. However, the broader adoption of IDMS in high-throughput studies remains limited by the high costs of commercially available ^13^C-labeled internal standards (IS), labor-intensive in-house IS production, and the narrow applicability of existing methods to different organisms. Here, we present a robust and scalable IDMS-based LC-MS/MS workflow for high-throughput metabolic profiling of diverse bacteria. The analytical method couples ion-pairing liquid chromatography with multiple reaction monitoring (MRM) to quantify 96 intracellular metabolites in under 16 minutes, with an average RSD of 21%. We developed a protocol for large-scale production of high-quality ^13^C-labeled IS, considerably lowering the cost for high-throughput IDMS studies. We then applied the workflow to 5 bacterial species in different cultivation conditions. This work provides a versatile platform for microbial metabolomics, supporting systems biology and data-driven metabolic engineering at scale.

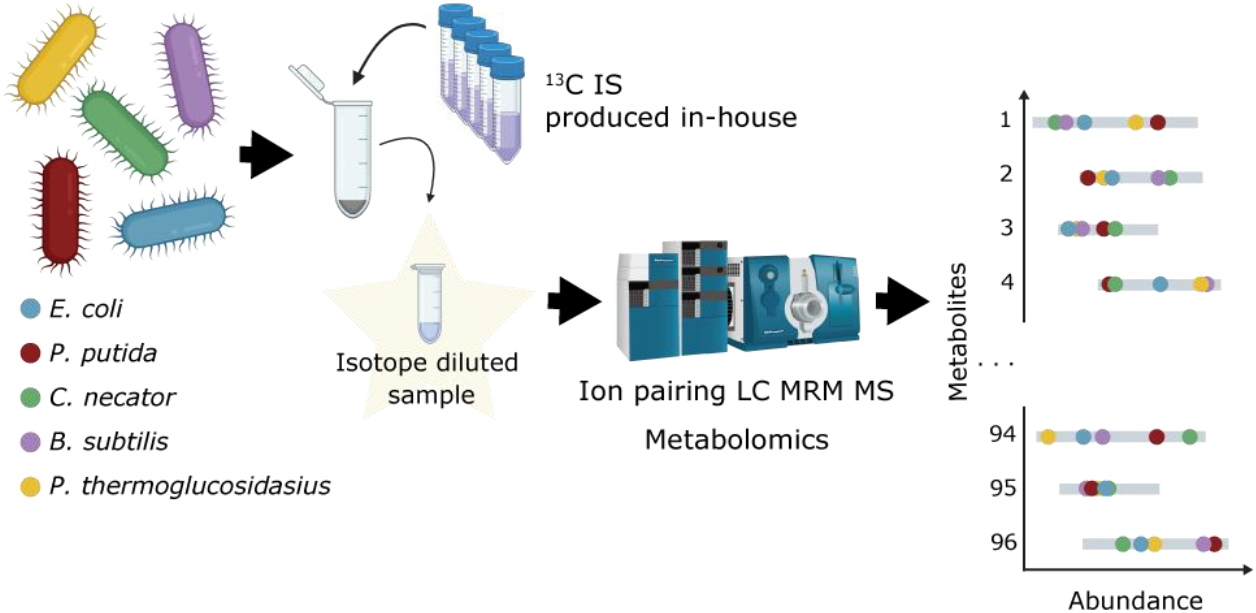

## INTRODUCTION

Quantitative metabolomics has become an essential tool for investigating the structure, function, and regulation of microbial metabolism (1,2). Over the past decades, stable isotope dilution mass spectrometry (IDMS) approaches have emerged as a gold standard for targeted metabolomics, offering enhanced accuracy in the quantification of intracellular metabolites (3– 6). When combined with multiple reaction monitoring (MRM), these methods can detect and quantify hundreds of intracellular metabolites within a few minutes (6–8), significantly improving analytical throughput. In parallel, filtration-based sampling techniques have emerged as a preferred approach for quenching cellular metabolism and preserving sample integrity, further improving the quality of metabolomic datasets (5,9). Accurate metabolite concentration measurements not only reveal the metabolic state of a cell (10,11) but are also essential for identifying signaling metabolites (12,13), understanding gene regulatory networks (14,15), and parameterizing kinetic models for model-guided metabolic engineering (16).

A central challenge in the broader application of IDMS is its dependency on isotopically labeled internal standards (IS). These are critical to compensate for variability inherent to LC-MS techniques caused by sample matrix effects and ion suppression during electrospray ionization (3,5,17). While commercially available ^13^C-labeled standards can be a practical solution, they offer a limited range of metabolites and are prohibitively expensive for large-scale applications. For example, using the Cambridge Isotope Laboratories, Inc. U-^13^C metabolite extracts of yeast (ISO1) or *Escherichia coli* (MSK-CRED-DD-KIT) adds approximately $20 or $30 per sample to the cost of an IDMS analysis. A widely adopted alternative approach is the in-house production of ^13^C-labeled cell extracts. This provides higher flexibility and quantification accuracy, by enabling IS generation from the organism of interest, ensuring a near-unit isotope ratio for each metabolite quantified. In-house production can be easily performed by cultivating microbes in uniformly labeled ^13^C carbon sources and the IS extracts processed in a similar way to the experimental samples. Although filtration prior to quenching improves IS quality by minimizing salt carryover and mimicking sample matrices (9,18,19), it is labor-intensive and difficult to scale, particularly when large volumes are required for high-throughput applications.

Another challenge to routine application of IDMS analytical methods is the tradeoff between robustness and throughput. Longer methods (17,18), methods optimized for specific metabolite classes (20), or those involving complex sample generation and derivatization (6) offer greater retention time stability and accuracy but are unsuitable for high-throughput applications. Conversely, faster, streamlined protocols optimized for specific organisms and experimental contexts (7,8) may lack reusability across diverse projects and are often not suitable for cross-species comparisons. These gaps present a critical bottleneck, especially as systems biology and artificial intelligence (AI)-driven approaches increasingly depend on large, high-quality omics datasets (21,22).

To address these challenges, we developed a robust and versatile IDMS-based metabolomics pipeline designed for the high-throughput quantification of primary metabolites in diverse bacterial species. Our approach includes a cost-effective protocol for the production of large volumes of high-quality ^13^C-labeled IS, alongside an optimized LC-MS/MS workflow employing ion-pairing chromatography and MRM detection. We validated this workflow with different gram-negative and gram-positive model organisms, showcasing its applicability, precision, and scalability for microbial systems biology and metabolic engineering applications.

## METHODS

### Chemicals and Materials

Uniformly labelled ^13^C glucose was purchased from Cambridge Isotope Laboratories^®^. The analytical standards used to prepare the calibration curves were purchased from Sigma Aldrich^®^. LC-MS grade water was purchased from Fisher Scientific^®^ and all other chemicals and carbon sources from Sigma Aldrich^®^ or VWR^®^. The PVDF 0.22 and 0.45 µm, 25 mm membrane filters were purchased form Millipore^®^. The 500 mL PVDF 0.45 µm, 75 mm bottle-top filtration system was purchased from VWR^®^.

### Strains, Media and Cultivations

The following bacterial strains were used in this work: *Escherichia coli* K-12 MG1655, *Pseudomonas putida* KT2440, *Cupriavidus necator* H16, *Bacillus subtilis* 168, and *Parageobacillus thermoglucosidasius* DSM2542. Cultivations were performed in M9 minimal media containing 42.2 mM Na2HPO4, 27.5 mM KH2PO4, 8.5 mM NaCl, 11.3 mM NH4Cl, 1 mM MgSO4, 0.1 mM CaCl2, 8.3 mM FeCl3, 1 mL/L SL7 trace element solution (DSMZ S4331). Media to cultivate *B. subtilis* was supplemented with 0.05 g/L of L-Tryptophan. Cells were cultivated with either glycolytic sole carbon sources - 11 mM glucose (*E. coli* and *P. putida*), 28 mM glucose (*B. subtilis* and *P. thermoglucosidasius*) or 11 mM fructose (*C. necator*) – or non-glycolytic sole carbon sources - 80 mM formate (*C. necator*), 60 mM acetate (*P. putida*), 83 mM acetate (*P. thermoglucosidasius*), 40 mM malate (*B. subtilis*) or 50 mM succinate (*E. coli*). Metabolically labeled internal standard (^13^C IS) was produced from *E. coli* grown on uniformly labelled ^13^C glucose. Cultures were started from overnight pre-cultures grown in the respective M9 medium, inoculated from single colony on LB plates. Cultivations were performed in 10 mL of media using Pioreactor™ small-scale bioreactors at 900 rpm, unless stated otherwise. Incubation temperatures were: 30°C for *P. putida* and *C. necator*, 37°C for *E. coli* and *B. subtilis*, and 60°C for *P. thermoglucosidasius*.

### Sampling and Intracellular Metabolite Extraction

Cultures were sampled during the exponential phase of growth. Biological samples for metabolomics analysis were prepared by fast filtration of 1 mL of culture under vacuum. The filter was submerged in 1 mL of cold 40:40:20 acetonitrile:methanol:water + 0.1% formic acid buffer (quenching buffer) (19) and incubated at -20°C for a minimum of 6 h to ensure all cells lysed. The extracts were centrifuged and concentrated by centrifugal evaporation as previously described (9). Dried samples were stored at -80°C and reconstituted in 90 µL of LC-MS water prior to analysis. To prepare ^13^C IS, 200 mL of culture was fast filtered under vacuum and 100 mL of quenching buffer was added to the filtration system. After an overnight incubation at -20°C the buffer was vacuum filtered, centrifuged and aliquoted into 2 mL tubes prior to concentration by centrifugal evaporation. Each aliquot was reconstituted in 20 µL of LC-MS water at a concentration 10 times higher than a regular culture sample (Figure 2f). Isotope diluted samples were prepared by adding 10 µL of ^13^C IS to the culture sample, ensuring a 1:1 ^12^C:^13^C isotope concentration ratio.

### Metabolome Quantification

Cell extracts were analyzed by LC-MS/MS. LC separation was performed on a SCIEX Exion LC (AB SCIEX^®^) using an Acquity UPLC HSS T3 100 mm x 2.1 mm x 1.8 µm (Waters^®^). Mobile phase A was water + 10 mM tributylamine + 10 mM acetic acid + 2% isopropanol + 5% methanol and B was isopropanol. A chromatographic gradient was defined as 0, 0, 0.5; 2.5, 0, 0.5; 4.3, 2, 0.5; 4.5, 6, 0.5; 5.5, 6, 0.5; 5.8, 11, 0.5; 6.5, 11, 0.5; 7.5, 28, 0.5; 8, 53, 0.2; 10.8, 53, 0.2; 11, 0, 0.2; 13, 0, 0.5; 16 stop [time (min), eluent B (vol%), flow rate (mL/min)]. Oven temperature was 40°C, autosampler temperature was 10°C and the injection volume was 6 µL. MS analysis was performed on a SCIEX TripleQuad 6500+ (AB SCIEX^®^) operated in negative mode and using multiple reaction monitoring (MRM) for quantification of defined transitions (Sup. table S1). Electrospray ionization parameters were defined as: source temperature of 400°C, electrospray voltage of -4500 V, curtain gas of 40 psi, CAD gas of 8 and gas 1 and 2 of 60 psi. Analyzer parameters were optimized for each metabolite by manual tuning using analytical standards. The mass calibration was done using the polypropylene glycol (PPG) standard.

Samples were acquired using the scheduled MRM algorithm and the extracted ion scans (XIC) were processed in SCIEX OS^®^ 3.4.5. Among the metabolites targeted were amino acids, sugar phosphates, nucleotides, redox cofactors, carboxylic acids and coenzyme A derivatives. Quantification was performed by IDMS using ^13^C IS. The abundance ratio ^12^C:^13^C was calculated for each metabolite. Calibration curves of standards spiked with ^13^C IS were run for the adenylate nucleotides: ATP, ADP, AMP. The linear regression was defined as the range of concentration that deviated less than 50% from the maximum slope. All calibration curves had a R^2^>0.99. Metabolite abundance and concentrations were normalized to the optical density at sampling time. The LOD was defined as 1e3 cps peak height and signal to noise as 10. QCs and carry-over checks were performed within each batch.

### Data Analysis

All statistical analysis and data representation were done in R (R Core Team 2024). The relative standard deviation (RSD) in percentage was calculated as 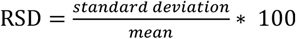. The adenylate energy charge ratio (AEC) was calculated as 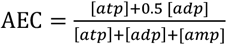.

## RESULTS and DISCUSSION

### Determination of filter pore size and buffer volume needed for effective sample filtration and cell lysis

Fast sampling by filtration enables the instantaneous quenching of cells, preventing the rapid metabolite turnover caused by exposing cells to stress or a different environment (23). The minimum volume of quenching buffer required for effective cell lysis during the cold incubation and the largest filter pore size capable of retaining cells were determined. To accelerate the filtration process, 0.45 µm filters were tested as a faster alternative to the 0.22 µm filters commonly used in the field (24–26). *E. coli* was cultivated with glucose as sole carbon source, and five 1 mL samples were taken (Figure 1a). Samples 1, 2, 3, and 4.1 were filtered through 0.45 µm filters and the filters were submerged in 100 µL, 250 µL, 500 µL, and 1 mL of quenching buffer, respectively. Sample 4.2 was filtered through a 0.22 µm filter and the filter was submerged in 1 mL of quenching buffer.

**Figure 1.**
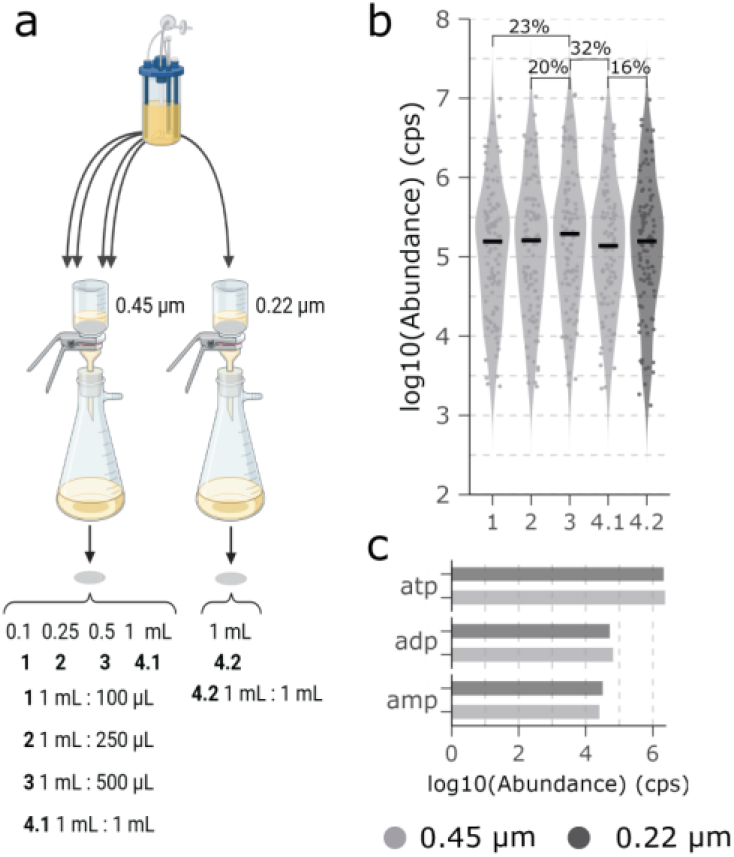
Comparison of filter pore size and buffer volume for sampling the intracellular metabolome of bacteria. **a** Schematic of the experimental workflow (created with BioRender.com). **b** Violin plots show the log-transformed median abundance of the 96 metabolites quantified. Percentage values indicate the median difference between selected samples. Colors indicate the filter pore size used. **c** Bar plot shows the log-transformed abundances of the adenylate metabolites with colors indicating filter pore size.

Median metabolite abundances across samples 1, 2, 3 and 4.1 differed by less than 32% (Figure 1b). Sample 3 showed the highest abundance, suggesting that a buffer-to-sample ratio of 1:2 may be optimal for effective lysis of cells on the filter. In samples 4.1 and 4.2, median metabolite abundances varied by 16.4% (Figure 1b). Moreover, the abundance of the adenylate metabolites, which have a fast turnover-time and are often used as an indication of effective sampling when calculating the energy charge ratio (10), varied by less than 20% (Figure 1c). These findings indicate that 0.45 µm filters retain cells nearly as effectively as 0.22 µm filters and that samples were quenched in comparable ways. Considering the cell sizes of the bacteria of interest (diameter: 0.5-1.2 µm; length: 0.9-7 µm) (27–31), this result was expected. Furthermore, filtration through 0.45 µm filters is approximately twice as fast as through 0.22 µm filters, considerably increasing the speed of sampling.

### Experimental workflow adjustments for in-house ^13^C IS production

To enable high-throughput isotope-ratio metabolomics, large volumes of ^13^C IS are required. Hence, a protocol to produce large volumes of ^13^C IS was developed. Sampling, filtration, concentration and processing were scaled and optimized to obtain a ^13^C IS of comparable quality to the samples produced by regular sampling workflow.

A PVDF 0.45 µm bottle-top vacuum filtration system was compared to PVDF 0.45 µm membrane filters, which are used for regular culture sampling. Filtering 1 mL of culture through a membrane filter takes *ca*. 20 s, whereas 200 mL through the filtration system takes ca. 3 min. Hence, it was essential to assess whether the filtration method and duration influenced the intracellular metabolome, and thereby the quality of the produced ^13^C IS. *E. coli* was cultivated in 1 L flasks with 200 mL of media, with U-^13^C glucose as sole carbon source. Sample 1 and 2 were filtered using the membrane filter and the filtration system, respectively (Figure 2a). The median metabolite abundance in sample 2 was 18.4% lower than in sample 1 (Figure 2b). Furthermore, the abundance of the adenylate metabolites differed by less than 20% between samples (Figure 2c), suggesting that the filtration system has a negligible impact on the extracted metabolome.

**Figure 2.**
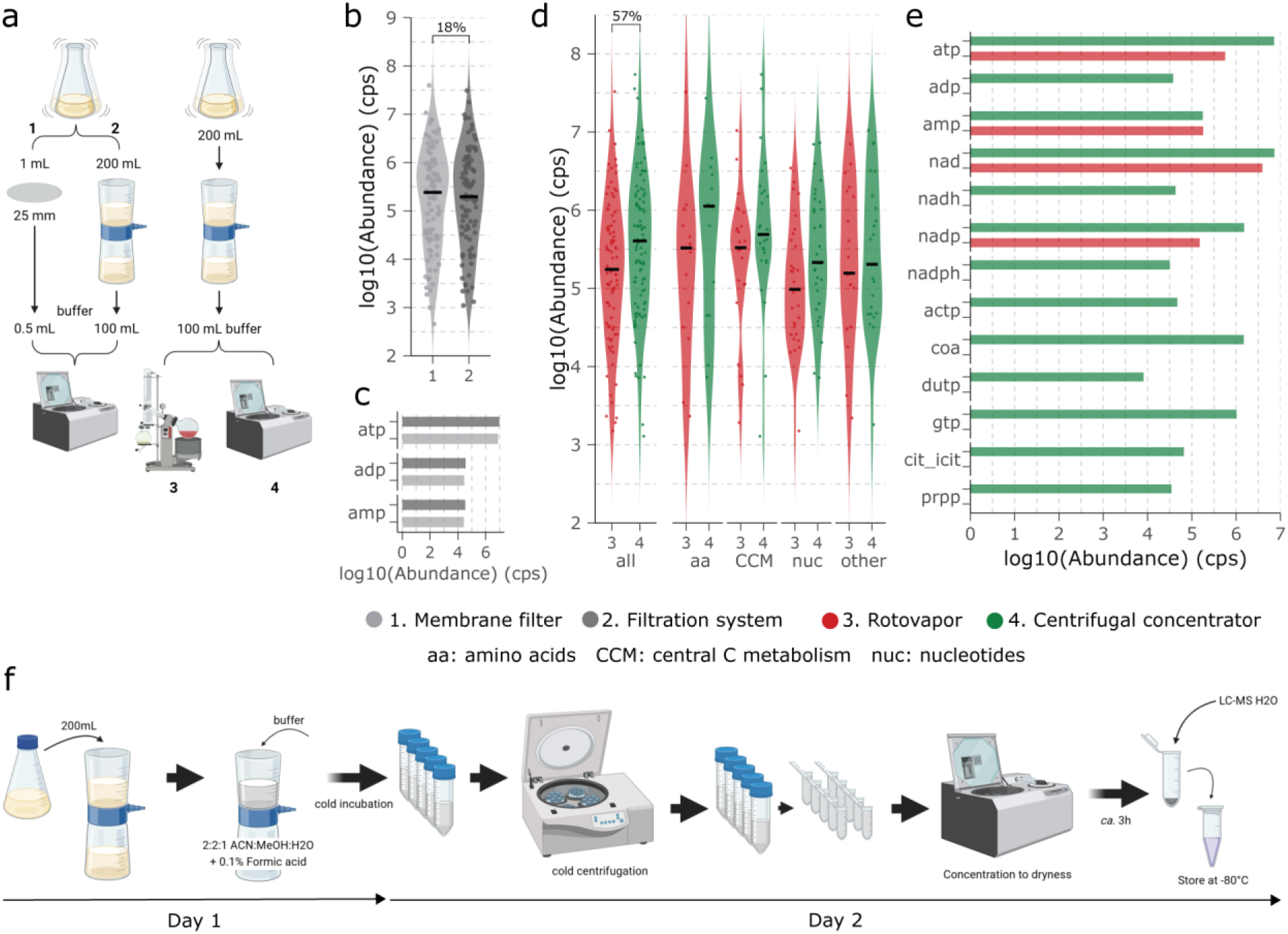
Sampling workflow adjustments for ^13^C IS production. **a** Schematic of the experimental workflow (created with BioRender.com). **b** Comparison of filtration systems. Violin plots show the log-transformed median abundance of the 96 metabolites quantified. Percentage value indicates the median difference between samples. Colors represent the filtration system used. **c** Bar plot shows the log-transformed abundance of the adenylate metabolites, with colors indicating the filtration method. **d** Comparison of concentration methods. Violin plots show the log-transformed median abundance: overall median of the 96 metabolites and medians by functional group. Percentage value indicates the median difference between samples. Colors represent the concentration method. **e** Bar plot shows the log-transformed abundance of selected metabolites, with colors indicating the concentration method. **f** Schematic of the experimental workflow for ^13^C IS production (created with BioRender.com).

To optimize sample concentration, rotary evaporation was compared to the usual centrifugal evaporation at 45°C. *E. coli* was cultivated in flasks with U-^13^C glucose as sole carbon source. Sample 3 and 4 were filtered through the filtration system, with sample 3 concentrated by rotary evaporation and sample 4 by centrifugal evaporation in 2 mL aliquots (Figure 2a). Metabolites were classified into four functional groups: amino acids, central carbon metabolism, nucleotides and others. The median metabolite abundance was calculated for both overall levels and individual groups. The median metabolite abundance of sample 4 is 57% higher than in sample 3 (Figure 2d), indicating more than 2-fold higher metabolite recovery when using centrifugal evaporation. Moreover, nine metabolites were undetectable in sample 3, most of which had abundances above 5×10^4^ cps in sample 4 (Figure 2e), further supporting the higher efficiency of centrifugal evaporation. Unlike centrifugal evaporation, rotary evaporation accelerates solvent removal through continuous rotation of the sample flask and increased surface area exposure. This leads to the dried extract spreading thinly along the flask walls, making complete re-dissolution difficult (32). Additionally, the faster and potentially harsher process may promote degradation of labile metabolites, such as redox cofactors, coenzymes and phosphorylated nucleotides. Although direct comparative studies involving rotary evaporation for sample preparation are limited, Bennour et al. reported reduced recovery of sensitive metabolites following rotary evaporation (32).

Following these results, a large amount of ^13^C IS was prepared as Sample 4 (Figure 2a). From 1 L of *E. coli* culture grown with 2 g of U-^13^C glucose, enough ^13^C IS was obtained to prepare 1000 isotope diluted samples (Figure 2f). Taking into consideration only material costs, our protocol enables the production of ^13^C IS in two days at a 20 times lower cost compared to commercially available alternatives.

### Robustness of the LC-MS method

To assess the robustness of the method, the RSD was calculated for all targeted metabolites in a biological sample injected 12 non-consecutive times over a 35-hour period. *E. coli* was cultivated with glucose as sole carbon source, and isotope diluted samples were analyzed. The RSD was calculated from the abundance ratio of all replicates. The average RSD was 21.3%, with 75% of metabolites showing an RSD below 30% (Figure 3, Figure S1a). This result suggests that, for most metabolites, an abundance ratio variation exceeding 30% is likely to reflect biologically meaningful differences between samples. Comparable RSD values have been reported in studies validating the robustness of similar LC-MS/MS methods for quantitative metabolic profiling (8,33,34).

**Figure 3.**
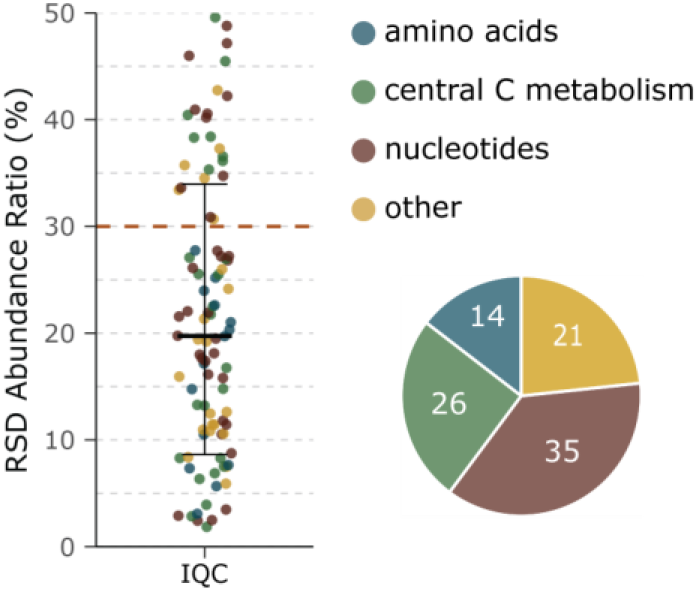
RSD as a measure of method robustness. Dots represent the % RSD across replicates (n=12) for the 96 metabolites quantified. The error bar represents the std deviation from the mean. The orange dashed line indicates the reference RSD threshold of 30%. The pie chart shows the number of metabolites quantified in each functional group.

To investigate factors contributing to measurement variability, parent ion mass, retention time (RT), and dwell time (DT) were correlated with the RSD (Figure S1b-d). RT and DT were the parameters most associated with variation in RSD (r^2^=0.52 and r^2^=0.31, respectively). Metabolites with RSD > 30% tended to elute later (after ∼6 min) and had lower DT (∼5 me) (Figure S1c,d). Notably, DT was correlated with RT (r^2^=0.5), suggesting that metabolites eluting later were quantified with lower reliability, likely due to shorter ion monitoring times during MRM acquisition (Figure S1e).

### Determination of the linear range of detection and application to various bacterial species

The linear range of detection for each metabolite was determined using a dilution series of a biological sample. *E. coli* was cultivated with glucose as sole carbon source. Sample X corresponds to 1 mL of culture, mimicking standard sampling volume. The additional samples 2X and 3X were obtained by sampling 2 mL and 3 mL, respectively. A 3-fold dilution series was prepared from sample X. Isotope diluted samples were prepared by adding 10 µL of ^13^C IS to each sample. The relative concentration range of the 96 metabolites quantified varied from 0.0014 to 3. Sample X was within the linear detection range for all metabolites, with 10.4% of metabolites quantified at the upper linear detection limit (Figure S2a).

To assess the applicability of the method across different bacterial species and conditions, gram-negative (*E. coli, C. necator, P. putida*) and gram-positive (*B. subtilis* and *P. thermoglucosidasius*) bacteria were cultivated in triplicate cultures on different media: *E. coli* on glucose or succinate, *C. necator* on fructose or formate, *B. subtilis* on glucose or malate, *P. putida* on glucose or acetate and *P. thermoglucosidasius* on glucose or acetate. Isotope diluted samples were analyzed. Approximately 83% of the abundance ratios fell within the respective linear detection ranges (Figure 4a). Most out-of-range measurements correspond to non-glycolytic growth conditions or gram-positive species, both of which were expected to exhibit more distinct metabolic profiles, considering the linear detection ranges were defined using extracts from *E. coli* grown on glucose. Nevertheless, the metabolic profiles across all conditions showed substantial overlap (Figure S2b), supporting the use of a ^13^C-labeled *E. coli* metabolome extract as a suitable IS for studying the metabolome of diverse microorganisms. For further analysis, measurements below the lower limit of the linear range were excluded and those above the upper limit of the linear range were capped at the maximum of the respective range.

**Figure 4.**
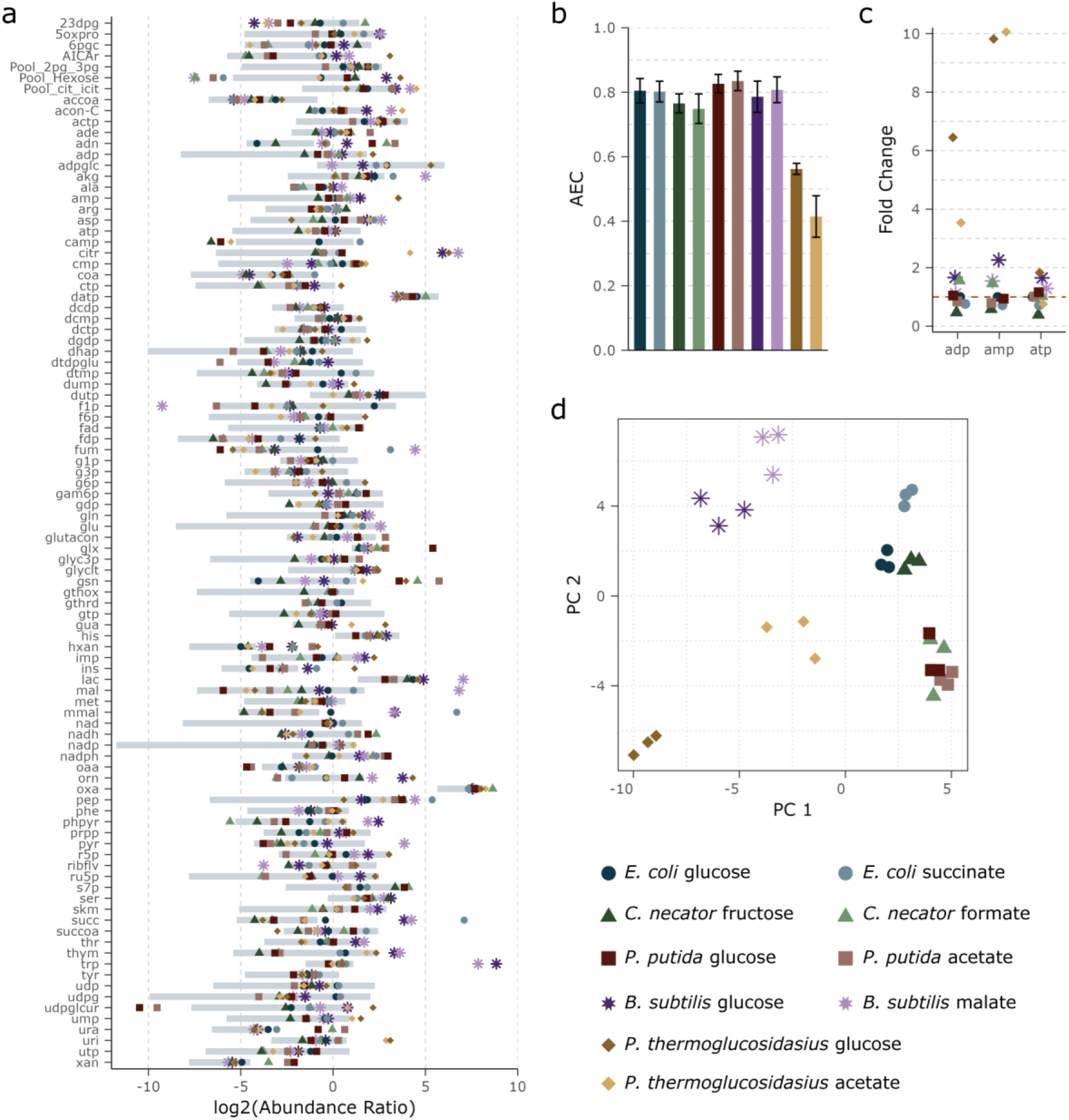
Linear range of detection and method application for metabolomics analysis of various bacteria. **a** Linear range of detection overlaid by measurements from various growth conditions for the 96 metabolites quantified. Bars represent the linear range expressed in log-transformed abundance ratio. Dots represent the log-transformed average abundance ratio (n=3) of different bacteria grown on different substrates. Symbols indicate the bacterial strain and colors the substrate. **b** Bar plot shows the adenylate energy charge ratio (AEC), with the error bars representing the std deviation from the mean (n=3). Colors represent the growth condition. **c** Scatter plot shows the fold change abundance ratio of the adenylate metabolites relative to E. coli cultivated in glucose. Symbols represent the bacterial strain and colors the substrate. **d** Principal component analysis (PCA) of metabolite abundance ratios, explaining 31.4% of the total variance (PC1:24.03%; PC2:17.38%). Symbols represent the bacterial strain and colors the substrate.

Adenylate nucleotide concentrations were quantified from 10-fold diluted samples. In most conditions, AEC values exceeded 0.75, indicating that cells remained metabolically active and maintained energy homeostasis across different substrates (Figure 4b). Notably, *P. thermoglucosidasius* showed considerably lower AEC values in both conditions, caused by elevated concentrations of ADP (3-6 times) and AMP (*ca*. 10 times), compared to other organisms, while ATP concentrations were comparable (Figure 4c). These results suggest that thermophilic bacteria might adopt different AECs than mesophilic bacteria. This was previously reported for *Bacillus stearothermophilus*, although with indirect measures of the adenylate pool and reporting also lower ATP content in thermophilic cells (35). An increased biosynthesis and pool of AMP and ADP in thermophiles might be necessary to maintain ATP homeostasis, by increasing ATP synthesis rates and preventing higher hydrolysis rates at higher temperatures (36). These higher pools of ADP might be compatible with the observation that different thermophiles evolved ADP-dependent kinases (37–39).

A principal component analysis (PCA) was performed to explore metabolic differences across growth conditions, revealing clear separation between Gram-positive and Gram-negative species, as well as additional species-specific clustering (Figure 4d). Gram-negative bacteria were characterized by elevated levels of energy cofactors and sugar phosphate intermediates, whereas Gram positive bacteria showed higher accumulation of nucleotide precursors and PPP intermediates (Figure S2c). Along PC2, the primary drivers of separation included TCA cycle intermediates and amino acids on the positive axis, and nucleotide related metabolites and redox cofactors on the negative axis (Figure S2d). Although less pronounced, glycolytic and non-glycolytic conditions were consistently separated within each bacterial species (Figure 4d), as expected (10). These findings highlight the strength of the method to capture both inter- and intra-strain metabolic variation.

## CONCLUSIONS

We present a robust and scalable isotope dilution LC-MS/MS workflow for the high-throughput quantification of intracellular metabolites across diverse bacterial species. The method demonstrates strong reproducibility for most metabolites, with an average RSD of 21% and over 75% of metabolites falling below the 30% RSD threshold, enabling confident interpretation of biological variation. A cost-effective and efficient filtration-based workflow for producing large volumes of high quality ^13^C-labeled IS was established and validated. Sampling and extraction procedures were optimized for high metabolite recovery and compatibility with high-throughput formats. The considerably lower production cost of the high quality ^13^C-labeled IS facilitates the broader application of IDMS methods in high-throughput studies. Importantly, the majority (∼83%) of metabolite measurements across species and carbon sources fell within their defined linear detection range, with dilution strategies enabling accurate quantification for remaining targets. The workflow allows the reliable separation of inter- and intra-species metabolic phenotypes and was used to highlight the considerably lower adenylate energy charge of a thermophilic bacterium compared to mesophilic bacteria. Together, these features establish our method as a powerful tool for microbial systems biology and large-scale metabolomics studies.

## ACKNOWLEDGMENTS

We thank Baris Kara and Belinda Escher for their help in setting up thermophilic cultivations. This work was supported by the Novo Nordisk Foundation (NNF20CC0035580 and NNF14OC0009473).

## AUTHOR CONTRIBUTIONS

**Catarina Rocha**: Conceptualization, Methodology, Formal Analysis, Investigation, Writing - Original Draft, Writing - Review & Editing, Visualization; **Sergi Pujol Pinto**: Formal Analysis, Investigation; **Sheila Ingemann Jensen**: Conceptualization, Resources; Lars Keld Nielsen: Conceptualization, Resources, Writing - Review & Editing, Supervision, Funding acquisition; **Stefano Donati**: Conceptualization, Methodology, Resources, Writing - Original Draft, Writing - Review & Editing, Supervision, Project Administration.

## Supplementary Materials

**Figure S1.**
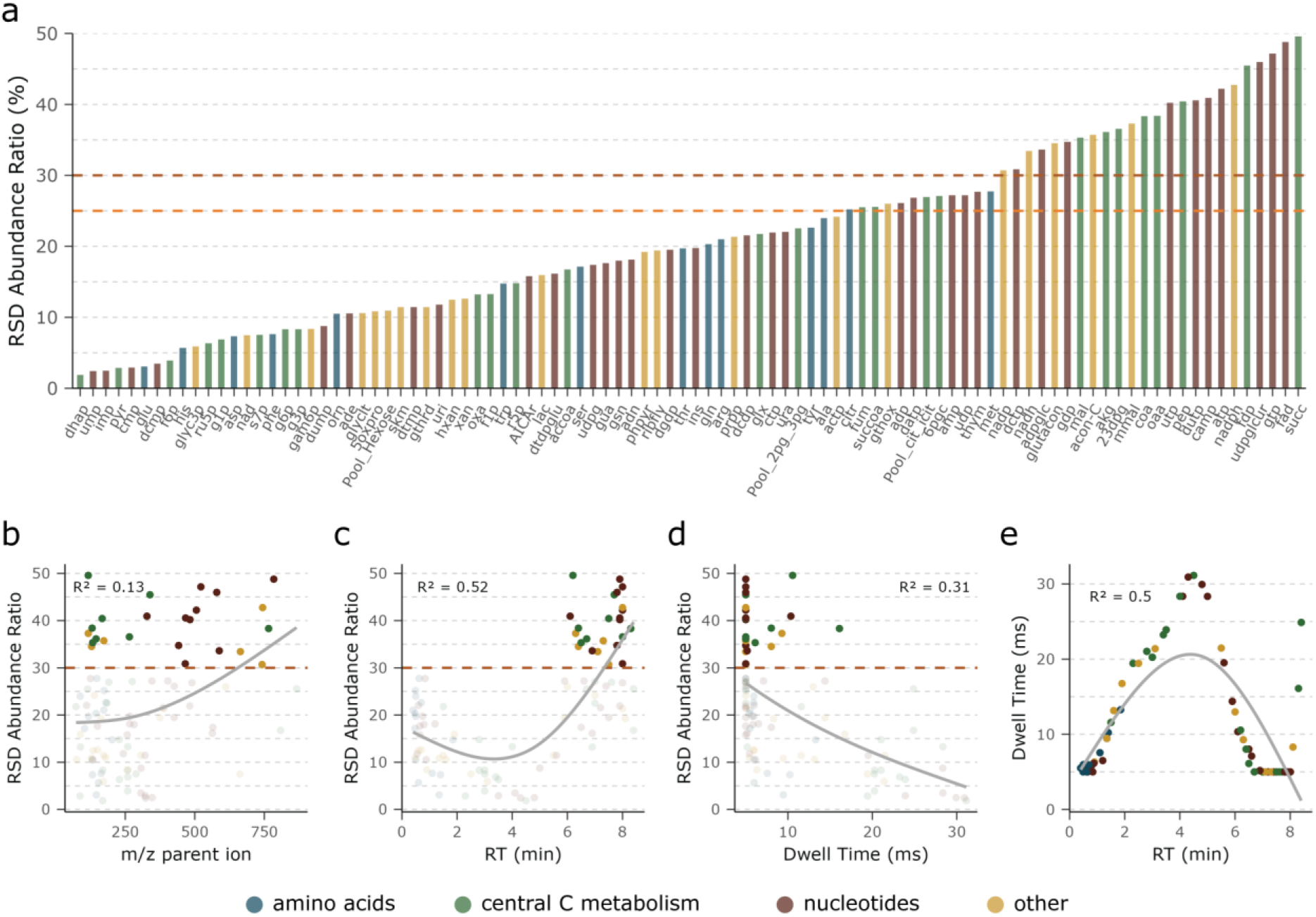
RSD as a measurement of method robustness (supplementary). **a** Bar plot shows the % RSD across replicates (n=12) for individual metabolites. The orange dashed lines indicate reference RSD thresholds of 25% and 30%. Colors represent metabolite functional groups. **b** Correlation plot between RSD and parent ion mas, with metabolites exceeding 30% RSD highlighted. **c** Correlation plot between RSD and retention time, with metabolites exceeding 30% RSD highlighted. **d** Correlation plot between RSD and dwell time, with metabolites exceeding 30% RSD highlighted. **e** Correlation plot between dwell time and retention time. Colors represent metabolite functional groups.

**Figure S2.**
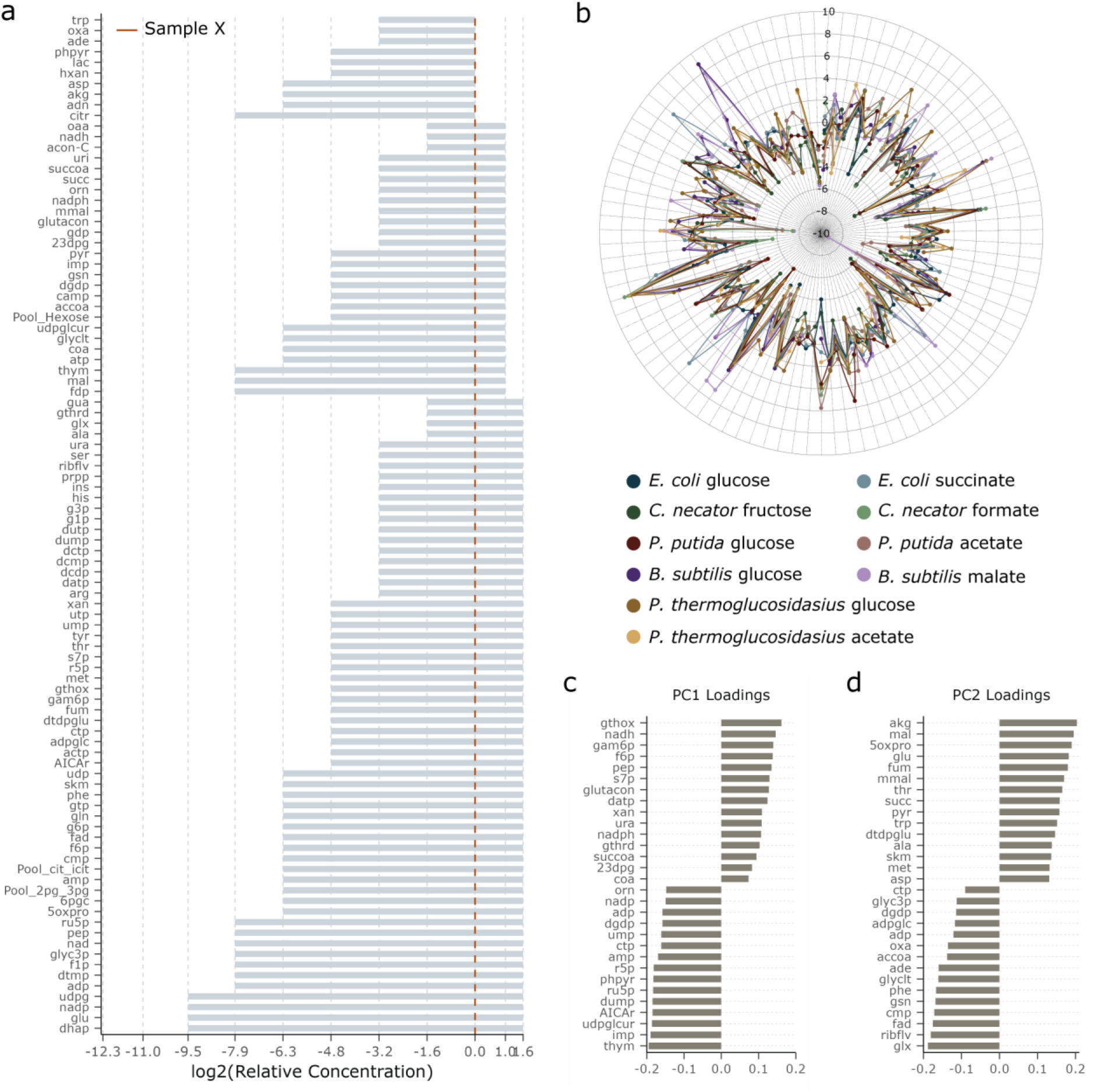
Linear range of detection and method application for metabolomics analysis of various bacteria (supplementary). **a** Linear range of detection for the 96 metabolites quantified. Bars represent the linear range expressed in log-transformed relative concentration. The orange dashed line indicates the reference sample at concentration 1. **b** Radial plot shows the overlaid metabolic profiles for all growth conditions. Dots represent the log-transformed abundance ratio. Colors indicate the growth condition. **c, d** Top 15 loadings for PC1 and PC2, ranked by their contribution to sample grouping.

## REFERENCES

1. Johnson CH, Ivanisevic J, Siuzdak G. Metabolomics: beyond biomarkers and towards mechanisms. Nat Rev Mol Cell Biol. 2016 July;17(7):451–9.

2. Donati S, Sander T, Link H. Crosstalk between transcription and metabolism: how much enzyme is enough for a cell? Wiley Interdisciplinary Reviews: Systems Biology and Medicine. 2018;10(1):e1396.

3. Mashego MR, Wu L, Van Dam JC, Ras C, Vinke JL, Van Winden WA, et al. MIRACLE: mass isotopomer ratio analysis of U-13C-labeled extracts. A new method for accurate quantification of changes in concentrations of intracellular metabolites. Biotechnology and Bioengineering. 2004;85(6):620–8.

4. Wu L, Mashego MR, van Dam JC, Proell AM, Vinke JL, Ras C, et al. Quantitative analysis of the microbial metabolome by isotope dilution mass spectrometry using uniformly 13C-labeled cell extracts as internal standards. Analytical Biochemistry. 2005 Jan 15;336(2):164–71.

5. Bennett BD, Yuan J, Kimball EH, Rabinowitz JD. Absolute quantitation of intracellular metabolite concentrations by an isotope ratio-based approach. Nat Protoc. 2008 Aug;3(8):1299–311.

6. Røst LM, Brekke Thorfinnsdottir L, Kumar K, Fuchino K, Eide Langørgen I, Bartosova Z, et al. Absolute Quantification of the Central Carbon Metabolome in Eight Commonly Applied Prokaryotic and Eukaryotic Model Systems. Metabolites. 2020 Feb;10(2):74.

7. McCloskey D, Xu J, Schrübbers L, Christensen HB, Herrgård MJ. RapidRIP quantifies the intracellular metabolome of 7 industrial strains of E. coli. Metabolic Engineering. 2018 May;47:383–92.

8. Guder JC, Schramm T, Sander T, Link H. Time-Optimized Isotope Ratio LC–MS/MS for High-Throughput Quantification of Primary Metabolites. Analytical Chemistry. 2017 Feb 7;89(3):1624–31.

9. McCloskey D, Utrilla J, Naviaux RK, Palsson BO, Feist AM. Fast Swinnex filtration (FSF): a fast and robust sampling and extraction method suitable for metabolomics analysis of cultures grown in complex media. Metabolomics. 2015 Feb 1;11(1):198–209.

10. Radoš D, Donati S, Lempp M, Rapp J, Link H. Homeostasis of the biosynthetic *E. coli* metabolome. iScience. 2022 July 15;25(7):104503.

11. Valgepea K, Lemgruber R de SP, Meaghan K, Palfreyman RW, Abdalla T, Heijstra BD, et al. Maintenance of ATP Homeostasis Triggers Metabolic Shifts in Gas-Fermenting Acetogens. cels. 2017 May 24;4(5):505–515.e5.

12. Lempp M, Farke N, Kuntz M, Freibert SA, Lill R, Link H. Systematic identification of metabolites controlling gene expression in *E. coli*. Nat Commun. 2019 Oct 2;10(1):4463.

13. Kochanowski K, Gerosa L, Brunner SF, Christodoulou D, Nikolaev YV, Sauer U. Few regulatory metabolites coordinate expression of central metabolic genes in Escherichia coli. Mol Syst Biol. 2017 Jan;13(1):903.

14. Mülleder M, Calvani E, Alam MT, Wang RK, Eckerstorfer F, Zelezniak A, et al. Functional Metabolomics Describes the Yeast Biosynthetic Regulome. Cell. 2016 Oct 6;167(2):553–565.e12.

15. Fuhrer T, Zampieri M, Sévin DC, Sauer U, Zamboni N. Genomewide landscape of gene–metabolome associations in Escherichia coli. Molecular Systems Biology. 2017;13(1):907.

16. Volkova S, Matos MRA, Mattanovich M, Marín de Mas I. Metabolic Modelling as a Framework for Metabolomics Data Integration and Analysis. Metabolites. 2020 Aug;10(8):303.

17. McCloskey D, Gangoiti JA, Palsson BO, Feist AM. A pH and solvent optimized reverse-phase ion-paring-LC– MS/MS method that leverages multiple scan-types for targeted absolute quantification of intracellular metabolites. Metabolomics. 2015 Oct 1;11(5):1338–50.

18. Buescher JM, Moco S, Sauer U, Zamboni N. Ultrahigh Performance Liquid Chromatography™Tandem Mass Spectrometry Method for Fast and Robust Quantification of Anionic and Aromatic Metabolites. Anal Chem. 2010 June 1;82(11):4403–12.

19. Rabinowitz JD, Kimball E. Acidic Acetonitrile for Cellular Metabolome Extraction from Escherichia coli. Anal Chem. 2007 Aug 1;79(16):6167–73.

20. Wahl SA, Seifar RM, ten Pierick A, Ras C, van Dam JC, Heijnen JJ, et al. Quantitative Metabolomics Using ID-MS. In: Krömer JO, Nielsen LK, Blank LM, editors. Metabolic Flux Analysis: Methods and Protocols [Internet]. New York, NY: Springer; 2014 [cited 2025 Aug 8]. p. 91–105. Available from: 10.1007/978-1-4939-1170-7_6

21. Noor E, Cherkaoui S, Sauer U. Biological insights through omics data integration. Current Opinion in Systems Biology. 2019 June;15:39–47.

22. Liebal UW, Phan ANT, Sudhakar M, Raman K, Blank LM. Machine Learning Applications for Mass Spectrometry-Based Metabolomics. Metabolites. 2020 June;10(6):243.

23. Taymaz-Nikerel H, De Mey M, Baart G, Maertens J, Heijnen JJ, van Gulik W. Changes in substrate availability in Escherichia coli lead to rapid metabolite, flux and growth rate responses. Metabolic Engineering. 2013 Mar 1;16:115–29.

24. Thorfinnsdottir LB, García-Calvo L, Bø GH, Bruheim P, Røst LM. Optimized Fast Filtration-Based Sampling and Extraction Enables Precise and Absolute Quantification of the Escherichia coli Central Carbon Metabolome. Metabolites. 2023 Jan 18;13(2):150.

25. Bolten CJ, Kiefer P, Letisse F, Portais JC, Wittmann C. Sampling for Metabolome Analysis of Microorganisms. Anal Chem. 2007 May 1;79(10):3843–9.

26. Bordag N, Janakiraman V, Nachtigall J, González Maldonado S, Bethan B, Laine JP, et al. Fast Filtration of Bacterial or Mammalian Suspension Cell Cultures for Optimal Metabolomics Results. PLoS One. 2016 July 20;11(7):e0159389.

27. Cullum J, Vicente M. Cell growth and length distribution in Escherichia coli. J Bacteriol. 1978 Apr;134(1):330–7.

28. Makkar NS, Casida LE. Cupriavidus necator gen. nov., sp. nov.; a Nonobligate Bacterial Predator of Bacteria in Soil†. International Journal of Systematic and Evolutionary Microbiology. 1987;37(4):323–6.

29. Nguyen CT, Nguyen THH, Tra VT, Tungtakanpoung D, Tran CS, Vo TKQ, et al. Paraquat removal by free and immobilized cells of Pseudomonas putida on corn cob biochar. Case Studies in Chemical and Environmental Engineering. 2023 Dec 1;8:100376.

30. Errington J, van der Aart LT. Microbe Profile: Bacillus subtilis: model organism for cellular development, and industrial workhorse. Microbiology (Reading). 2020 May;166(5):425–7.

31. Najar IN, Thakur N. A systematic review of the genera Geobacillus and Parageobacillus: their evolution, current taxonomic status and major applications. Microbiology. 2020 Sept 1;166(9):800–16.

32. Bennour N, Mighri H, Eljani H, Zammouri T, Akrout A. Effect of solvent evaporation method on phenolic compounds and the antioxidant activity of Moringa oleifera cultivated in Southern Tunisia. South African Journal of Botany. 2020 Mar;129:181–90.

33. Pezzatti J, Bergé M, Boccard J, Codesido S, Gagnebin Y, H Viollier P, et al. Choosing an Optimal Sample Preparation in Caulobacter crescentus for Untargeted Metabolomics Approaches. Metabolites. 2019 Oct;9(10):193.

34. Takenaka M, Yoshida T, Hori Y, Bamba T, Mochizuki M, Vavricka CJ, et al. An ion-pair free LC-MS/MS method for quantitative metabolite profiling of microbial bioproduction systems. Talanta. 2021 Jan;222:121625.

35. Webster JJ, Walker BG, Leach FR. ATP content and adenylate energy charge ofBacillus stearothermophilus during growth. Current Microbiology. 1988 Sept 1;16(5):271–5.

36. Moeller C, Schmidt C, Guyot F, Wilke M. Hydrolysis rate constants of ATP determined in situ at elevated temperatures. Biophysical Chemistry. 2022 Nov;290:106878.

37. Tuininga JE, Verhees CH, Oost J van der, Kengen SWM, Stams AJM, Vos WM de. Molecular and Biochemical Characterization of the ADP-dependent Phosphofructokinase from the Hyperthermophilic Archaeon Pyrococcus furiosus *. Journal of Biological Chemistry. 1999 July 23;274(30):21023–8.

38. Kengen SWM, Tuininga JE, de Bok FAM, Stams AJM, de Vos WM. Purification and Characterization of a Novel ADP-dependent Glucokinase from the Hyperthermophilic Archaeon Pyrococcus furiosus(*). Journal of Biological Chemistry. 1995 Dec 22;270(51):30453–7.

39. Hansen T, Schönheit P. ADP-dependent 6-phosphofructokinase, an extremely thermophilic, non-allosteric enzyme from the hyperthermophilic, sulfate-reducing archaeon Archaeoglobus fulgidus strain 7324. Extremophiles. 2004 Feb 1;8(1):29–35.

